# Enrichment Probe Sets Combining Universal and Lineage-Specific Targets Help Resolve Recalcitrant Lineages

**DOI:** 10.64898/2026.03.24.713849

**Authors:** Irene Villa-Machío, Irene Masa-Iranzo, Nicolai M. Nürk, Lisa Pokorny, Andrea S. Meseguer

## Abstract

The combination of target capture sequencing (TCS) with low-coverage whole genome sequencing (lcWGS), an approach known as Hyb-Seq, has allowed the integration of natural history collections into the genomics revolution, transforming biodiversity research. To implement Hyb-Seq, a collection of genomic targets, often nuclear orthologs, is needed to design probes for TCS. In flowering plants, the universal Angiosperms353 probe set has been proven resolutive at multiple evolutionary scales, with caveats. Malpighiales is known to be one of the most challenging flowering plant orders to resolve. Within this order, the clusioid clade (∼2.2K species, 94 genera, five families) is no exception. To resolve phylogenetic relationships in this recalcitrant clade, we design a custom probe set, the Clusioids626 kit, composed of 39,936 120-mer probes targeting 626 nuclear orthologs (∼6.6M nucleotides). This probe set includes all Angiosperms353 targets and 273 clusioid-specific ones, carefully chosen taking copy-number, length evenness, and phylo-informativeness into account. We test our probe set on 70 accessions representing all families and tribes in the clusioid clade. On average, 50.4% of TCS reads mapped to our targets, recovering a median of ∼600 orthologs. Relationships for all clusioid families are fully resolved for our nuclear targets. Additionally, 105 plastid coding DNA sequences were retrieved from the lcWGS fraction. A strong cyto-nuclear conflict was detected. The Clusioids626 kit performs better than the universal Angiosperms353 enrichment panel alone. Our kit design workflow can be extended into other lineages for which a universal probe set exists but more resolution is needed.

## 1 Introduction

Target capture sequencing (TCS) has emerged as a standard phylogenomic approach in systematics and evolution (McKain et al. 2018; Andermann et al. 2020). In particular, TCS combined with low-coverage whole genome sequencing (lcWGS), an approach known as Hyb-Seq (Weitemier et al. 2014; Dodsworth et al. 2019), can result in hundreds to thousands of nuclear genomic loci (e.g., nuclear orthologs), as well as high-copy number DNA (e.g., organellar DNA), useful for phylogenomic inference in non-model organisms (Breinholt et al. 2021; Ametrano et al. 2025). This technique has recently become very popular, as it has been shown to be both cost efficient (Hale et al. 2020) and effective resolving phylogenetic relationships for flowering plants across evolutionary scales (from micro to macroevolutionary levels; Villaverde et al. 2018; Zuntini et al. 2024) and for DNA templates of varying quality, including degraded herbarium material (Brewer et al. 2019; Shee et al. 2020). The wealth of molecular data generated by this technique has been proven useful to generate robust phylogenetic hypotheses and to resolve species radiations (Kadlec et al. 2017; Mitchell et al. 2017; Larridon et al. 2020), but also to investigate gene tree conflict and the causes leading to reticulation, be them methodological (e.g., gene tree estimation error) or biological, i.e., incomplete lineage sorting (ILS), horizontal gene transfer, hybridization, etc. (Fleming et al. 2023; Steenwyk et al. 2023; Bjornson et al. 2024).

Gene capture techniques have followed two main strategies. One approach relies on universal hybridization capture kits designed to target large clades across the tree of life (Buddenhagen et al. 2016; Breinholt et al. 2021; Waycott et al. 2021). The most popular in plants being the Angiosperms353 kit designed to target all flowering plants (Johnson et al. 2019). This universal kit resolves deep phylogenetic relationships across distantly related taxa, and has also demonstrated utility in resolving shallow relationships (Larridon et al. 2020; Slimp et al. 2021; Beck et al. 2021; Balant et al. 2025), though their effectiveness at finer scales remains debated for certain lineages (Yardeni et al. 2022). This has led to the development of customized probe sets tailored to specific plant lineages. To date, custom kits are available for numerous plant families, such as Euphorbiaceae (Villaverde et al. 2018), Dioscoreaceae (Soto Gomez et al. 2019), Ochnaceae (Schneider et al. 2021), Scrophulariaceae (Villaverde et al. 2023) or Asteraceae (Moore-Pollard et al. 2024), among others. More recently, efforts to combine universal and lineage-specific probe sets into a single kit have shown great promise (Eserman et al. 2021; Ogutcen et al. 2021; Fonseca et al. 2023, 2024; Bentz and Leebens-Mack 2024; Musker et al. 2025). These probe sets integrate the broad applicability of universal kits with the high resolution of specific ones, providing an efficient and versatile framework for phylogenomic studies (Fonseca et al. 2024). However, the use of kits combining universal and specific probe sets is still quite limited, in spite of their potential.

The order Malpighiales accounts for ∼7.8% of eudicot diversity and includes over 16K species, across ∼700 genera and 36 families (APG IV 2016; Zuntini et al. 2024). This order has been referred to as one of the most recalcitrant angiosperm groups, since resolving phylogenetic relationships at various levels has proven to be quite the ordeal (Wurdack and Davis 2009). Phylogenetic relationships in this group remain poorly understood, even with inferences based on ∼100 genes from three genomic compartments (Xi et al. 2012). This difficulty has been attributed to its rapid radiation in the Mid-Cretaceous (Davis et al. 2005; Xi et al. 2012) and the challenges of phylogenetic inference, referred to as a ‘perfect storm’ of ILS, introgression, and gene tree estimation error that blurs the resolution of Malpighiales, particularly with regards to the backbone topology (Cai et al. 2021). Within order Malpighiales, the clusioid clade comprises ∼2.2K species, 94 genera, and five families: Bonnetiaceae, Calophyllaceae, Clusiaceae, Hypericaceae, and Podostemaceae (Wurdack and Davis 2009; Ruhfel et al. 2011; Xi et al. 2012). The clusioid clade is nearly cosmopolitan, although the majority of its species are found in the tropics, in a broad range of habitats—closed canopy rainforests, dry, temperate, and high-altitude tropical forests, and swift-flowing aquatic environments—and forms, e.g., trees, shrubs, and herbs (Stevens 2007; Ruhfel et al. 2016; Meseguer et al. 2018). The terrestrial members of the clade are key components of tropical rainforest ecosystems (Davis et al. 2005); while Podostemaceae, as the largest strictly aquatic plant family of the clade, plays an essential role in riverine systems (Philbrick and Novelo 1995; Cook 1996). Several species of the clusioid clade are economically important, them being edible (e.g., the mangosteen, *Garcinia mangostana* L., and the mammee apple, *Mammea americana* L.) or used for timber production (*Calophyllum brasiliense* Cambess., *Mesua ferrea* L.), pharmaceutical purposes (*Hypericum perforatum* L.), or in horticulture (several Calophyllaceae, Clusiaceae, and Hypericaceae species) (Ernst 2003; Stevens 2007; Ruhfel et al. 2011).

Relationships within the clusioid clade, based on morphological data, were so unclear that the existence of the group itself was contested. Families Calophyllaceae and Hypericaceae were nested within Clusiaceae *s.l.*, also known as Guttiferae (e.g., Robson 1977; Notis 2004), and some Podostemaceae genera were placed in distantly related orders, such as Theaceae (Ericales, Wawra 1886; Cronquist 1981; Takhtajan 1997), making the placement of Podostemaceae uncertain and speculative. In the last decades, several molecular studies have sought to disentangle relationships of Malpighiales and the clusioid clade (Korotkova et al. 2009; Wurdack and Davis 2009; Ruhfel et al. 2011; Jin et al. 2020; Cai et al. 2021) and within clusioid families (Calophyllaceae: Trad et al. 2021; Clusiaceae: Gustafsson et al. 2002; Marinho et al. 2019; Gaudeul et al. 2024; Hypericaceae: Meseguer et al. 2013, 2015; Nürk et al. 2013, 2015; Podostemaceae: Tippery et al. 2011; Bedoya et al. 2019). However, despite the advances recently made, reconstructing phylogenetic relationships within the clusioid clade remains challenging, with the topology changing depending on the molecular markers used or the analytical approach implemented.

Wurdack and Davis (2009), relying on 13 gene regions from all three plant genomic compartments, confirmed the monophyly of the clusioids and recovered a Bonnetiaceae plus Clusiaceae clade, sister to a clade composed of Calophyllaceae sister to Hypericaceae plus Podostemaceae. Ruhfel et al. (2011) obtained the same topology using three plastid and one mitochondrial genes, with the most extensive sampling to date (194 spp.). So did Xi et al. (2012) using genes from all three genomic compartments, and Jin et al. (2020) using 82 protein-coding plastid genes. Based on plastomes, Trad et al. (2021) mostly supported the topology of Wurdack and Davis (2009), but highlighted major gene tree discordance, e.g., at the divergence between Bonnetiaceae and Clusiaceae. Recently, Cai et al. (2021) presented a different topology based on 423 nuclear genes, and showed extensive gene tree discordance. This latter topology did not include Podostemaceae and showed Clusiaceae successively sister to Hypericaceae, and a Bonnetiaceae plus Calophyllaceae clade. In general, relationships within the Bonnetiaceae, Clusiaceae, and Podostemaceae families are largely consistent across different studies, but this is not the case for all other clusioid families (Ruhfel et al. 2011; Trad et al. 2021).

In this study, we design a new custom probe set, the Clusioids626 kit, which combines nuclear ortholog genes specific to the clusioid clade, with genes from the universal Angiosperms353 enrichment panel. Additionally, we develop a specific plastid target file for the clusioid clade to recover plastid genes from Hyb-Seq off-target reads. The Clusioids626 kit, together with the plastid target file, was employed to infer phylogenetic relationships within the clusioid clade. We compare the performance of our Clusioids626 probe set against that of the Angiosperms353 universal panel. Finally, we also analyze gene tree conflict within and across genomic compartments.

## 2 Materials and Methods

### 2.1 Taxon Sampling for Custom Probe Set Design

To design the Clusioids626 probe set we used 12 transcriptomes, one belonging to Ochnaceae (putatively sister to the clusioid clade; Zuntini et al. 2024) and 11 belonging to the five clusioid families: five Hypericaceae, three Calophyllaceae, two Clusiaceae, and one Bonnetiaceae (Table S1). Transcriptomic data were downloaded from the sequence read archive (SRA) database, hosted by the National Center for Biotechnology Information (NCBI), and were assembled de novo using Trinity v.2.9.1 (Grabherr et al. 2011) from quality-filtered reads obtained using Trimmomatic v.0.36 (Bolger et al. 2014).

### 2.2 Probe Set Design and Loci Discovery

Two pipelines were used to select and mine target loci: MarkerMiner v.1.0 (Chamala et al. 2015) and HybPiper v.2.1.6 (Johnson et al. 2016). First, we identified putative single-copy nuclear orthologs using MarkerMiner (with default settings). This pipeline relies on the curated list of 2,809 mostly single-copy genes (De Smet et al. 2013) from the PLAZA Comparative Genomics Platform (Van Bel et al. 2012). As genomic inputs, we used the 12 transcriptome assemblies listed above and *Ricinus communis* L. (Euphorbiaceae, Malpighiales; v.0.1; Chan et al. 2010) as reference genome, available from Phytozome (Goodstein et al. 2012). As a result, a total of 1,637 nuclear orthologs shared across clusioids were identified (Table S2). Subsequently, we examined whether our list of candidate gene sequences overlapped with those included in the universal Angiosperms353 enrichment panel (Johnson et al. 2019) using BLAST+ (Camacho et al. 2009). Out of the 353 single-copy targets of the Angiosperms353 enrichment panel, 217 were not recovered by MarkerMiner. Second, we used the HybPiper pipeline, which requires a reference (i.e., a fasta file with the intended targets), to mine the remaining 217 targets, which we got from the mega353.fasta target file (McLay et al. 2021; available from github.com/chrisjackson-pellicle/NewTargets). We generated an ad hoc clusioid target file that included all the gene sequences extracted from MarkerMiner, plus the gene sequences from the mega353.fasta target file. The 12 transcriptomes were then mined with this ad hoc clusioid target file to obtain matrices with sequence information for each of the selected target genes. These target-gene data matrices were aligned with MAFFT v.7.520 (Katoh and Standley 2013), using default settings, and summary statistics were calculated with AMAS v.1.0 (Borowiec 2016). For probe design, final loci were selected that met the following criteria: *(i)* all loci contained sequences of at least seven species; *(ii)* three of them belonged to genus *Hypericum*, the most species-rich genus in the clusioid clade (WFO 2024); with *(iii)* minimum target length of 350 bp; and *(iv)* proportion of phylo-informative sites per target gene of at least 0.08. These filters resulted in a final set of 626 targets.

Biotinylated RNA probes were designed by Daicel Arbor Biosciences (Ann Arbor, MI, USA). Malpighiales Phytozome genomes *Linum usitatissimum* L. v.1.0, *Manihot esculenta* Crantz v.8.1, *Populus trichocarpa* Torr. & A.Gray ex Hook. v.4.1, *Ricinus communis* v.0.1, and *Salix purpurea* L. v.5.1, together with NCBI genomes *Bruguiera* Lam., *Hevea* Aubl., *Hypericum*, and *Viola* L., were used to remap the probes and to check that they did not overlap with high-copy-number elements, organellar DNA inclusive.

The final set had 39,9K 120-mer probes with 1x tiling density. Given this tiling density and to ascertain the efficacy of the probes in recovering our clusioid targets, BLAST+ was run on the probes against said target sequences. Various parameters were calculated, including the percent identity (i.e., percentage of nucleotides that are the same between the two sequences), alignment length, and percent coverage. The mean percent coverage of the probes for each target was extracted using the command-line program GNU datamash (FSF 2014).

### 2.3 Taxon Sampling, Genomic Library Preparation, and Probe Set Validation

To validate our probe set, we used 70 samples, 67 belonging to seven families of Malpighiales (Table S3) and three to order Celastrales. Out of those 70 samples, 38 were processed in-house with the Clusioids626 kit (see molecular workflow below) and 32 were downloaded from the European Nucleotide Archive (ENA) and mined for our 626 targets (see bioinformatic pipeline below). The clusioid clade was represented by 58 species from its five families: 19 Calophyllaceae, 19 Hypericaceae, seven Podostemaceae, ten Clusiaceae, and three Bonnetiaceae (Table S3). Species from other putatively closely related Malpighiales families were included as outgroups (six Ochnaceae, three Linaceae; Table S3). Species names follow WFO (2024).

Genomic DNA was extracted from 38 herbarium collections (Table S3) following the CTAB protocol (Doyle and Doyle 1987) with modifications (Larridon et al. 2020; Shee et al. 2020). All DNA extracts were quantified using a Qubit 3 Fluorometer with dsDNA high sensitivity (HS) assay kits (ThermoFisher Scientific, Waltham, MA, USA). Genomic DNA with fragment sizes ≥ 500 bp were sonicated to an average fragment size of 250 bp using a Covaris M220 Focused-ultrasonicator (Woburn, MA, USA).

Libraries were prepared using the DNA NEBNext Ultra II Library Prep Kit and Dual Index Primers (Multiplex Oligos) Sets 1 and 2 for Illumina^®^ (New England BioLabs, Ipswich, MA, USA), following the manufacturer’s protocol, but adjusting for half volume reactions (Hale et al. 2020). After end-repair, adapter ligation, and cleaning, libraries were indexed (10-cycle polymerase chain reaction, PCR), and then cleaned with AMPure XP magnetic beads (Beckman Coulter Life Sciences, Indianapolis, IN, USA). The samples were then quality-checked with a Bioanalyzer High Sensitivity DNA chip on a 2100 Bioanalyzer (Agilent Technologies, Santa Clara, CA, USA) and quantified using the Qubit as above. Genomic libraries were multiplexed in equimolar pools for target capture.

Pools were enriched using the Clusioids626 kit following the manufacturer’s myBaits® Manual v.5.02. Hybridizations were performed at 65°C for all genomic libraries. The enriched pools were amplified for 12, 14, or 16 PCR cycles (depending on DNA template quality) using the primers P5 and P7 for Illumina^®^ (Meyer and Kircher 2010) and PCR products were cleaned with AMPure XP beads. Final products were again run on the Bioanalyzer as above and then they were pooled equimolarly for sequencing. After multiplexing 75% enriched library pools with 25% plain genomic libraries, for all species, sequencing was performed on an Illumina^®^ HiSeq (San Diego, CA, USA) at Macrogen (Seoul, South Korea), producing 2 × 150 bp long reads.

### 2.4 Quality Filtering of Raw Reads and Sequence Assembly

Demultiplexed reads were inspected before and after trimming with FastQC 0.12.1 (Andrews 2010) for quality control. Raw reads were paired, adapter cleaned, and trimmed using fastp v.0.23.4 (Chen et al. 2018) with settings: -f 15, -t 10, -F 15, -T 15, -g, -W 4, -r, -M 20, -q 15, -l 50. Sequences were recovered using HybPiper with our Clusioids626 target file as the reference. The trimmed and paired reads were initially mapped against the target loci using BWA v.0.7.17 (Li and Durbin 2009), followed by de novo assembly using SPAdes v.3.15.5 (Bankevich et al. 2012), with default parameters. HybPiper assembly was repeated for samples that recovered less than 100 genes with sequences > 50% of the mean target length, lowering the coverage cutoff down to 3. When gene recovery was low, samples were rerun using BLASTx (Camacho et al. 2009) instead of BWA. The presence of potential paralogs was investigated using the paralog_retriever parameter and potential paralogs were removed if they had > 2 sequences/gene in ≥ 2 samples.

### 2.5 Data Matrix Construction and Multiple Sequence Alignment (MSA)

We retrieved FASTA files (HybPiper retrieve_sequences script) containing the exon-only sequences for 70 samples to build the corresponding nuclear data matrices. To identify and discard suboptimal samples and/or regions, and to build balanced data matrices (Shee et al. 2020), we implemented in R v.4.2.1 (R Core Team 2024) the max_overlap script (available from github.com/keblat/bioinfo-utils/blob/master/docs/advice/scripts/max_overlap.R). We filtered out underrepresented, incomplete, and unevenly distributed sequences and species (coverage score < ⅔ median value) to reduce the noise in our matrices.

Sequence matrices were aligned in MAFFT using default settings. We computed the summary statistics of the individual MSAs with AMAS and removed genes with values < ⅓ of the median for alignment size and proportion of Parsimony-informative characters (P_PIC_). Exploratory gene trees were generated using FastTree v.2.1.11 (Price et al. 2010) and analyzed in TreeShrink v.1.3.7 (Mai and Mirarab 2018) with the ‘per-species’ option, to automatically remove outlier taxa by excluding branches that increased the diameter of each gene tree beyond a given threshold (α parameter). Summary statistics were calculated with AMAS for all data matrices, so that we could choose the α parameter (0.50 threshold) that resulted in the highest proportion of P_PIC_, the lowest proportion of missing data, and the smallest data matrix size. Each locus was then realigned using MAFFT with default settings. We further refined the MSAs by trimming each gene using trimAl v.1.4.1 (Capella-Gutiérrez et al. 2009) with relaxed settings (-gt 0.3 -cons 30) and, once more, summarised the MSAs stats with AMAS. We removed genes whose alignment size and P_PIC_ median values were < ⅓.

### 2.6 Phylogenomic Analyses

To reconstruct phylogenetic relationships in the clusioid clade, we implemented both multispecies coalescent (MSC) and maximum likelihood estimation (MLE) approaches.

#### 2.6.1 Nuclear tree inference

Shrunk and trimmed MSAs were employed to generate MLE gene trees using IQ-TREE2 v.2.2.3 (Minh et al. 2020). The best substitution model for each gene was selected according to ModelFinder Plus (MFP; Kalyaanamoorthy et al. 2017). The resulting collection of MLE unrooted gene trees was used to infer the species tree with ASTRAL-III v.5.7.8 (Zhang et al. 2018) under the MSC approach. Before running ASTRAL-III, branches in gene trees with low support were collapsed using the nw_ed function from the Newick Utilities v.1.6 toolkit (Junier and Zdobnov 2010) to improve the accuracy of species tree inference (Mirarab 2019; Simmons and Gatesy 2021). Branch support in ASTRAL-III was calculated as local posterior probabilities (LPPs), and gene congruence was evaluated using normalised quartet scores (QS) values. These scores, which range from 0 to 1, represent the proportion of input gene tree quartets that are consistent with the species tree (the higher the score, the less discordant the gene trees are). We implemented the *astralProjection* function, available from the *AstralPlane* package (Hutter 2021) in R to plot pie charts depicting these QS values. An additional nuclear MLE analysis was conducted in IQ-TREE2 by concatenating all shrunk and trimmed MSAs into a supermatrix with AMAS. Genes were partitioned and allowed to evolve at independent rates and under different substitution models (determined by MFP). Support for the concatenated nuclear MLE tree was calculated (1,000 replicates) using ultrafast bootstrap (UFB; Hoang et al. 2018) and SH-like approximate likelihood ratio tests (SH-aLRT; Anisimova et al. 2011). All phylogenetic trees were visualized and plotted with FigTree v.1.4.4 (Rambaut 2018).

#### 2.6.2 Amino Acid (AA) Plastid Target File and Organellar Tree Inference

We mined plastid coding DNA sequences (CDS) from quality-checked paired reads of 70 clusioid clade samples and related taxa using HybPiper and a custom-designed AA plastid target file comprising 105 CDS. The stepwise design of this AA plastid target file started with the complete plastome sequences (downloaded from NCBI) of 14 species belonging to all clusioid families (Table S4). Following a first-pass mining of our 70 samples in HybPiper, we added the sequences of 13 more samples into our final AA plastid target file. These 13 samples were selected to reduce phylogenetic biases, that is, samples were chosen to maximize phylogenetic spread. The recovered CDS were concatenated into a single data matrix and a MLE plastid tree was inferred using IQ-TREE2, partitioned by CDS, and MFP was used to select the best substitution model for each CDS. Support was calculated using UFB and SH-aLRT (1,000 replicates).

### 2.7 Probe Sets Comparison: Clusioids626 vs. Angiosperms353

To evaluate the performance of our custom Clusioids626 probe set against the universal Angiosperms353 enrichment panel, we compared HybPiper summary statistics, as well as phylogenetic support and resolution of nuclear phylogenies inferred for data matrices mined with both our nuclear Clusioids626 and mega353.fasta target files for all 70 samples. Nuclear matrices containing exon-only sequences were shrunk and filtered following the same procedure described above.

### 2.8 Assessing Gene- vs. Species-Tree and Cyto-Nuclear Conflict

We assessed topological discordance and conflict between the MSC nuclear and MLE plastid ultrametric topologies (obtained in R with the *chronos* function from the *ape* package; Paradis and Schliep 2019) using the *tanglegram* and *untangle* functions (*step1side* method) from the *dendextend* package (Galili 2015) in R.

For the nuclear dataset, we also performed network analyses to explore potential gene trees vs. species tree conflicts and to detect patterns of reticulation caused by hybridization and/or ILS. The concatenated-genes matrix, with 552 loci and a selection of 58 samples (including only representatives of the five clusioid families), was imported into SplitsTree v.6.3.35 (Huson and Bryant 2024) using the neighbor-net algorithm with uncorrected P-distances to compute a split network (1,000 bootstrap replicates). To further explore the contribution of hybridization in the evolutionary history of the clusioid clade, we inferred phylogenetic networks using PhyloNet v.3.8.2 (Wen et al. 2018), which enables the inference of reticulate nodes (hybridization events). Due to computational limitations, for this analysis we reduced the taxon sampling to 12 representative species (two species from each clusioid family, and two outgroups), and used a set of 63 nuclear gene trees that could be rooted using Celastraceae as the outgroup. These nuclear trees were generated with RAxML-NG v.1.2.0 (Kozlov et al. 2019) and rooted with *Stackhousia minima* Hook.f. (Celastrales) with the *pxrr* command in *Phyx* (Brown et al. 2017). Phylogenetic networks, allowing for one to four reticulations, were inferred under maximum pseudo-likelihood, and five optimal phylogenetic networks were returned for each run. The command *CalGTProb* was used to compute the likelihood scores of the best network, scores that were then compared using a LRT. All phylogenetic networks were visualized using Dendroscope3 (Huson and Scornavacca 2012).

## 3 Results

### 3.1 The Clusioids626 Probe Set

Our nuclear data matrix includes 5,290 target sequences, for a total size of 6,568,704 nucleotides, resulting in 39,936 probes padding 626 target genes. The median alignment length for these genes is 1,742 bp (± 1,023.82 SD) and a median P_PIC_ of 0.14 (Table S5). The biotinylated probes have a mean alignment length of 111.43 bp, mean identity percentage to the target data matrix of 96%, and mean coverage of the probes per target of 93.50%. The Clusioids626 kit targets three distinct sets of genes: (i) 273 nuclear genes singled out by MarkerMiner and not present among the Angiosperms353 targets; (ii) 150 Angiosperms353 targets not found by MarkerMiner; and (iii) 203 custom-selected genes singled out by MarkerMiner and present in the Angiosperms353 panel.

### 3.2 Target Enrichment and Data Mining Performance

We produced novel TCS data for 36 species belonging to the clusioid clade plus one outgroup (*Linum suffruticosum* L.) that, together with the SRA data downloaded from NCBI, amounted for a total of 70 accessions. On average, 24.3 million paired reads were generated per sample and kept after quality filtering (ranging from 0.64 to 539.71 million reads; see heatmaps in Figure S1).

The percentage of on-target reads varied across specimens, ranging from 0.20% to 86.10% (mean = 31%, median = 29.40%; Table 1, Figure S2), and families. Hypericaceae has the highest mean percentage of on-target reads (56.34%), followed by Calophyllaceae (39.55%). Podostemaceae exhibits the lowest on-target read-recovery rate (2%; Table 1, Figure S2). In total, 608 of 626 targeted loci (97.12%) were recovered across the 70 specimens analyzed (Figure S1, Table S6). A median of 481.5 loci were recovered at a 50% target length (mean = 395.30, min = 30, max = 608). The capture success rate was measured as the percentage of the total captured length of all target loci per individual relative to the total mean length of all loci. Please, note that total captured length may exceed target length, meaning percentages can surpass 100%. The mean capture success rate was 110.40% (Table S6), ranging from 13.22% (*Garcinia portoricensis*, Clusiaceae) to 201.70% (*Hypericum tomentosum,* Hypericaceae).

**Table 1.** Summary statistics of target capture sequencing (TCS) success, including total number of filtered raw reads (NumReads), percent of on-target reads (PctOnTarget), the number of genes mapping to target (GenesMapped), and the number of loci retained at 50% target length (GenesAt50pct).

| Family | Taxon | Name | NumReads | PctOnTarget | GenesMapped | GenesAt50pct |
| --- | --- | --- | --- | --- | --- | --- |
| Hypericaceae | *Harungana madagascarensis* | C105 | 5104596 | 56.3 | 626 | 601 |
| Hypericaceae | *Psorospermum senegalense* | C109 | 42102790 | 74.1 | 626 | 585 |
| Calophyllaceae | *Lebrunia bushaie* | C1102 | 641146 | 51.1 | 626 | 318 |
| Bonnetiaceae | *Ploiarium elegans* | C1103 | 13482066 | 58.4 | 626 | 458 |
| Hypericaceae | *Hypericum foliosum* | C114 | 11182924 | 82.3 | 626 | 607 |
| Calophyllaceae | *Haploclathra paniculata* | C1141 | 6794246 | 52.2 | 626 | 526 |
| Calophyllaceae | *Mesua ferrea* | C1142 | 32875932 | 82.9 | 626 | 557 |
| Calophyllaceae | *Kayea nuda* | C1143 | 7752542 | 14.4 | 626 | 507 |
| Calophyllaceae | *Clusiella elegans* | C1172 | 694940 | 24.1 | 626 | 98 |
| Calophyllaceae | *Poeciloneuron indicum* | C1174 | 18264412 | 35.5 | 626 | 598 |
| Hypericaceae | *Hypericum galioides* | C133 | 20169360 | 66.6 | 626 | 530 |
| Hypericaceae | *Eliea articulata* | C189 | 4899152 | 58.2 | 626 | 604 |
| Hypericaceae | *Harungana rubescens* | C192 | 3883046 | 54.1 | 626 | 598 |
| Hypericaceae | *Hypericum longistylum* | C301 | 8564486 | 86.1 | 626 | 607 |
| Hypericaceae | *Hypericum calcicola* | C366 | 3425804 | 73.6 | 626 | 608 |
| Hypericaceae | *Hypericum tomentosum* | C428 | 8813024 | 75.5 | 626 | 608 |
| Hypericaceae | *Hypericum elodes* | C69 | 9890866 | 43.8 | 626 | 529 |
| Hypericaceae | *Cratoxylum arborescens* | C702 | 4865406 | 53.2 | 626 | 589 |
| Hypericaceae | *Cratoxylum formosum subsp. formosum* | C704 | 3398946 | 42.6 | 626 | 581 |
| Hypericaceae | *Cratoxylum glaucum* | C705 | 4791328 | 61.2 | 626 | 591 |
| Hypericaceae | *Cratoxylum formosum subsp. pruniflorum* | C706 | 3756756 | 43.7 | 626 | 586 |
| Hypericaceae | *Vismia baccifera* | C708 | 2114140 | 42.2 | 626 | 540 |
| Hypericaceae | *Vismia bilbergiana* | C709 | 4022110 | 45.9 | 626 | 575 |
| Hypericaceae | *Vismia cayennensis* | C710 | 2536426 | 40.1 | 626 | 557 |
| Hypericaceae | *Vismia guianensis* | C711 | 3682204 | 44.6 | 626 | 582 |
| Calophyllaceae | *Calophyllum brasiliense* | C835 | 16456140 | 40.9 | 626 | 591 |
| Clusiaceae | *Garcinia brasiliensis* | C837 | 1268102 | 14.5 | 626 | 194 |
| Calophyllaceae | *Endodesmia calophylloides* | C839 | 9633928 | 74.8 | 626 | 552 |
| Calophyllaceae | *Caraipa llanorum* | C841 | 7003404 | 73.6 | 626 | 507 |
| Clusiaceae | *Symphonia globulifera* | C846 | 4007402 | 18 | 626 | 259 |
| Bonnetiaceae | *Bonnetia sessilis* | C848 | 2618940 | 4.5 | 626 | 92 |
| Clusiaceae | *Moronobea riparia* | C849 | 5665994 | 49.3 | 626 | 199 |
| Calophyllaceae | *Kielmeyera coriacea* | C851 | 4479618 | 76.6 | 626 | 458 |
| Calophyllaceae | *Mammea vatoensis* | C852 | 4457186 | 62.5 | 626 | 554 |
| Calophyllaceae | *Marila laxiflora* | C853 | 6811474 | 35 | 626 | 597 |
| Calophyllaceae | *Neotatea longifolia* | C854 | 16508018 | 32.4 | 626 | 597 |
| Linaceae | *Linum suffruticosum* | C947 | 2609744 | 20 | 625 | 200 |
| Podostemaceae | *Dalzellia ubonensis* | DRR238799 | 51026638 | 2.9 | 607 | 522 |
| Podostemaceae | *Hydrobryum japonicum* | DRR238809 | 40302216 | 1.4 | 590 | 486 |
| Podostemaceae | *Polypleurum stylosum* | DRR238819 | 36182322 | 3.9 | 588 | 477 |
| Podostemaceae | *Terniopsis brevis* | DRR238825 | 101763126 | 2.2 | 618 | 537 |
| Podostemaceae | *Weddellina squamulosa* | DRR238827 | 42205222 | 2.2 | 610 | 516 |
| Podostemaceae | *Zeylanidium tailichenoides* | DRR258785 | 84168300 | 0.3 | 576 | 203 |
| Linaceae | *Linum tenue* | ERR10035165 | 17142584 | 4.1 | 624 | 250 |
| Clusiaceae | *Tovomita fructipendula* | ERR10978005 | 1255758 | 41 | 586 | 93 |
| Calophyllaceae | *Mammea americana* | ERR2040359 | 16211672 | 2.9 | 625 | 324 |
| Clusiaceae | *Garcinia oblongifolia* | ERR2040362 | 16269602 | 4.2 | 621 | 418 |
| Clusiaceae | *Garcinia livingstonei* | ERR2040363 | 27122088 | 1.6 | 617 | 339 |
| Ochnaceae | *Ochna serrulata* | ERR2040388 | 23604924 | 2.4 | 607 | 380 |
| Ochnaceae | *Cespedesia spathulata* | ERR5034039 | 3339896 | 7.1 | 611 | 152 |
| Ochnaceae | *Godoya obovata* | ERR5034041 | 2562200 | 7.5 | 603 | 152 |
| Ochnaceae | *Brackenridgea zanguebarica* | ERR5034068 | 3332182 | 9.3 | 619 | 129 |
| Ochnaceae | *Campylospermum reticulatum* | ERR5034069 | 1699686 | 3.4 | 594 | 115 |
| Ochnaceae | *Fleurydora felicis* | ERR5034071 | 2445734 | 8.8 | 603 | 141 |
| Celastraceae | *Elaeodendron orientale* | ERR7618728 | 5193870 | 8.5 | 562 | 130 |
| Celastraceae | *Stackhousia minima* | ERR7619745 | 6571210 | 6.7 | 597 | 94 |
| Hypericaceae | *Cratoxylum cochinchinense* | ERR7621010 | 6267036 | 26.4 | 624 | 180 |
| Calophyllaceae | *Kielmeyera petiolaris* | ERR7621021 | 1898258 | 37.1 | 609 | 162 |
| Calophyllaceae | *Caraipa densifolia* | ERR7621022 | 2802468 | 35.3 | 614 | 131 |
| Clusiaceae | *Moronobea coccinea* | ERR7621023 | 5410120 | 36.9 | 605 | 175 |
| Celastraceae | *Tripterygium wilfordii* | SRR12059219 | 62865170 | 2.6 | 625 | 420 |
| Podostemaceae | *Marathrum utile* | SRR12956182 | 539716186 | 1.3 | 616 | 503 |
| Clusiaceae | *Garcinia anomala* | SRR14356990 | 75830742 | 0.2 | 625 | 213 |
| Calophyllaceae | *Calophyllum longifolium* | SRR15016251 | 19854228 | 3.6 | 626 | 504 |
| Bonnetiaceae | *Bonnetia paniculata* | SRR16214477 | 46248790 | 0.2 | 626 | 46 |
| Calophyllaceae | *Calophyllum inophyllum* | SRR17138441 | 41340556 | 0.4 | 626 | 30 |
| Clusiaceae | *Garcinia portoricensis* | SRR19237490 | 49098458 | 4.3 | 626 | 555 |
| Linaceae | *Linum usitatissimum* | SRR20318611 | 46938580 | 2.2 | 622 | 340 |
| Calophyllaceae | *Calophyllum macrocarpum* | SRR7412626 | 3849732 | 16.2 | 625 | 242 |
| Clusiaceae | *Clusia rosea* | SRR7412628 | 7447036 | 7.1 | 626 | 372 |
| **MEDIAN** |  |  |  | 29.40 | 626 | 481.50 |
| **MEAN** |  |  |  | 30.84 | 618.77 | 395.30 |

The median taxon occupancy per target locus was 56 species, with a median alignment length of 1,860 bp (±1,465.65 SD) and 0.41 P_PIC_ (Table S7), more than double the P_PIC_ value for the probe set target sequences alone (P_PIC_ = 0.14; Table S5). Nineteen genes were discarded because their coverage scores (max_overlap script) were < ⅔ of the median, six of these were Angiosperms353 targets. Additionally, six target loci were flagged for potential paralogs (≥ 3 variants), three were Angiosperms353 targets, and were excluded from downstream analyses. After outlier removal and matrix refinement, 552 nuclear MSAs were generated, with a median alignment length of 1271.50 bp (447–6,500 bp) and a P_PIC_ of 0.49 (0.27–0.66). The concatenated nuclear data matrix had a size of 767,230 bp.

### 3.3 Probe Sets Comparison

When processing our dataset with the Angiosperms353 target file, HybPiper recovered 343 genes (97.16% of targets) at a 50% target length across the 70 samples analyzed, with a median of 198 loci (mean = 190, min = 1, max = 343; Figure S1, Table S8). The percentage of on-target reads ranged from 0 to 18.5%, with a median value of 5.35% (mean = 4.85%) and with median taxon occupancy per locus of 56 taxa. MSAs lengths ranged from 192 to 4,892 bp, with a mean length of 956.96 bp (Tables 2, S9). In contrast, when processing our dataset with the Clusioids626 target file, the MSA lengths ranged from 351 to 12,092 bp, with an average MSA length of 2,135.78 bp (Tables 2, S7).

Regardless of the target file used to process the data, the larger the MSA length the greater the number of Parsimony-informative characters (PICs); however, the highest PIC values were obtained for data processed with the Clusioids626 target file (Figure S3). Regarding taxon occupancy (Table 2), it was higher for the Clusioids626 target file (mean = 55.67 taxa, min = 36, max = 69) than for the Angiosperms353 target file (mean = 44.87 taxa, min = 13, max = 67). The percent of capture success was highest for genes mined with the Clusioids626 target file (110.40%), recovering an average of 483.56 loci (77.25% of all Clusioids626 targets; Table S6); whereas, for data generated with the Angiosperms353 target file, capture success was lower (51.68%) and we recovered an average of 226.61 loci (64.12% of all Angiosperms353 targets; Table S6).

**Table 2.** Number of taxa, length of the aligned contigs, percent of missing data, and number of Parsimony-informative characters (P_PIC_) for two different datasets: A) aligned contigs of the loci targeted with the Clusioids626 probe set, and B) aligned contigs of the loci targeted with the Angiosperms353 enrichment panel.

|  |  | No. of taxa | Alignment length | Missing percent | Parsimony informative sites |
| --- | --- | --- | --- | --- | --- |
| **A) Clusioid626 (608 genes)** | **Mean** | 55.67 | 2135.78 | 45.54 | 776.82 |
|  | **SD** | 6.49 | 1465.65 | 15.09 | 401.73 |
|  | **Max** | 69 | 12092 | 83.6 | 3097 |
|  | **Min** | 36 | 351 | 4.41 | 160 |
|  | **Total** | 70 | 1298556 |  | 472309 |
| **B) Angiosperms353 (343 genes)** | **Mean** | 44.87 | 956.96 | 27.63 | 423.90 |
|  | **SD** | 18.70 | 651.19 | 14.08 | 254.89 |
|  | **Max** | 67 | 4892 | 75.73 | 1450 |
|  | **Min** | 13 | 192 | 1.69 | 89 |
|  | **Total** | 70 | 330151 |  | 146246 |

We evaluated topological support, resolution, and discordance for the MSC nuclear trees generated with sequences assembled either with the Clusioids626 or the Angiosperms353 target files. For each analysis, we calculated normalized quartet scores (QS), the effective number of genes per branch/node (EN), the total number of quartets in all the gene trees supporting the main topology (t1), and local posterior probabilities (LPPs; Table S10). In the Clusioids626 MSC nuclear phylogeny, the proportion of gene trees per node supporting the primary topologies (t1) exceeded ⅓ for 67 nodes, 0.50 for 52 nodes, and 0.75 for 35 nodes (Figure S4). Similarly, Angiosperms353 t1 showed a gene tree proportion > ⅓ for 64 nodes, 0.50 for 49 nodes, and 0.75 for 35 nodes (Figure S4). In both cases, the mean proportion of gene trees per node supporting t1 was 0.73. The Clusioids626 MSC phylogeny showed 45 nodes, out of 66 nodes, with maximum support (LPP = 1), 17 with high support (1.0 > LPP ≥ 0.9), three nodes with moderate support (0.9 > LPP ≥ 0.7), and two of them with low support (LPP < 0.7). On the other hand, the Angiosperms353 MSC tree showed 40 out of 64 nodes with maximum support, 17 with high support, five with moderate support, and five with low support (Table S10).

### 3.4 Plastid Data Matrix Assembly

We successfully recovered sequences from 105 plastid loci from off-target reads (Table S11), achieving a mean taxon occupancy of 39 taxa, a mean alignment length of 1,328 bp (90–22,892 bp), and a P_PIC_ of 0.24 (Table S12). Following the calculation of coverage scores with the max_overlap script, 14 taxa were excluded from the plastid dataset (median coverage score < ⅓). Similarly, 11 plastid CDS were discarded (median coverage score < ⅔). Additionally, five plastid regions were flagged by HybPiper as having potential paralogs (≥ 3 variants), and were excluded from downstream analysis. Once outlier removal and matrix optimization were completed, MSAs were inferred for 64 plastid CDSs. These MSAs had a median alignment length of 591 bp (170–5,759 bp) and a P_PIC_ of 0.24 (0.10–0.71). The total alignment length of the concatenated plastid data matrix is 60,498 bp.

### 3.5 Phylogenomic Inference

Nuclear species trees were strongly supported for both MSC and MLE analyses (Figures 1, S5, and S6). Both the MSC and MLE nuclear topologies support the monophyly of the clusioid clade and its five constituent families. Trees were rooted with Celastrales. The MSC nuclear species tree was inferred from 552 MLE gene trees, with bipartitions collapsed for SH-aLRT support values = 0 (collapsing under various bootstrap thresholds did not alter the species tree topology but affected support values). Navigating the topology from the tips towards the root, the nuclear phylogeny recovered a Podostemaceae plus Hypericaceae clade sister to Calophyllaceae, with these three families sister to Bonnetiaceae, and all of them sister to Clusiaceae (LPP = 1). Comparisons of quartet frequencies supporting t1 versus the two alternative topologies (t2 and t3) showed minimal gene tree discordance, with QS = 0.92 (Figure 1). The MLE nuclear tree derived from the analysis of the concatenated dataset had maximal UFB and SH-aLRT support values (100%) across most branches (Figure S6). This concatenated MLE tree recovered the same topology as the MSC nuclear species tree. A nuclear topology from data mined using the Angiosperms353 target file was inferred under the MSC framework. This MSC Angiosperms353 tree recovered the same overall topology as the MLE and MSC nuclear trees obtained from the Clusioids626 targets. However, some unsupported variation was observed within family Calophyllaceae, especially pertaining genera *Clusiella*, *Marila*, and *Neotatea* (Figure S7).

**Figure 1.**
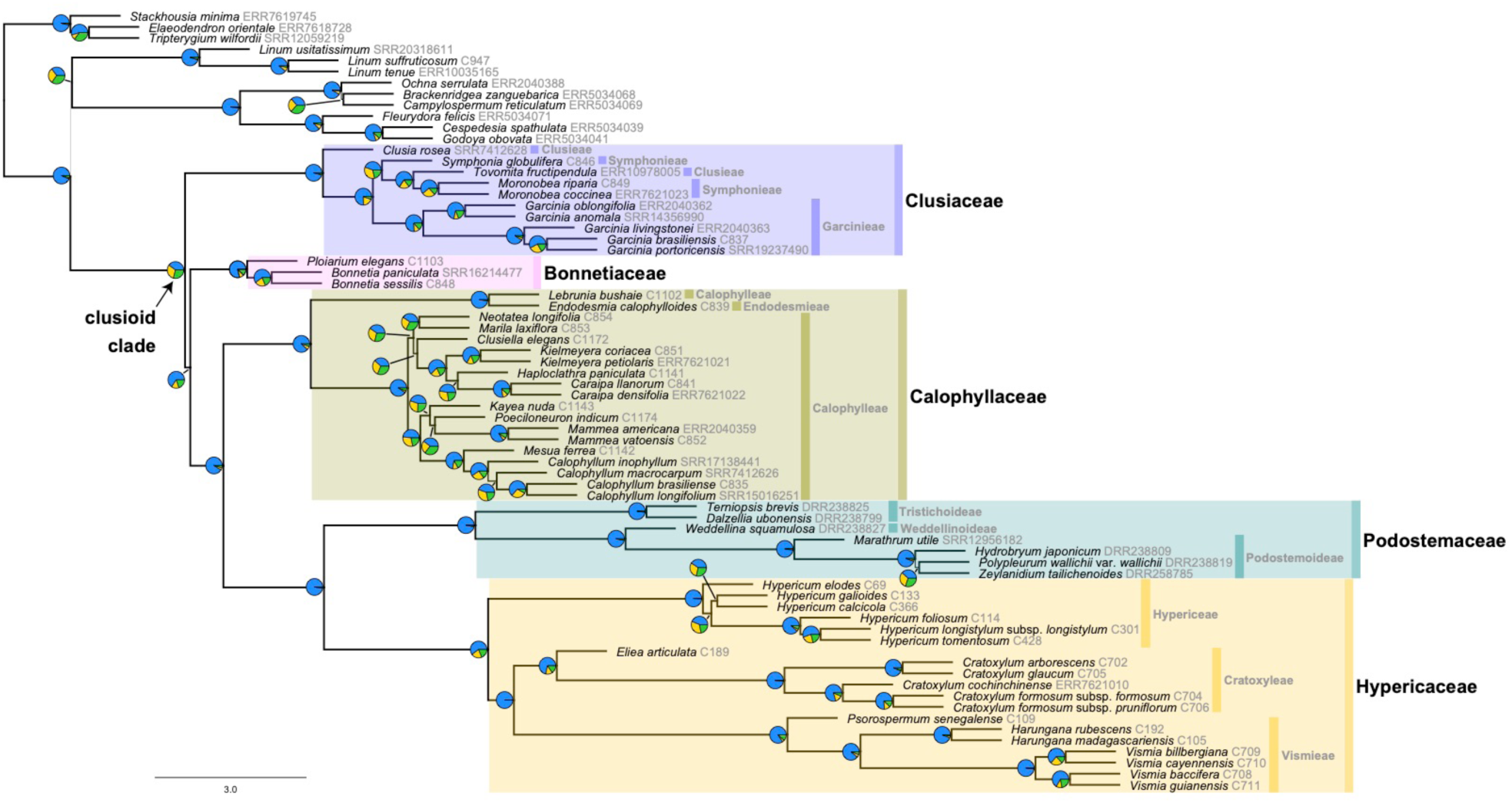
Nuclear species tree inferred under the MSC in ASTRAL-III, for 70 clusioid-clade and related taxa, from 552 targets assembled with our Clusioids626 target file. The thickness of the branches is proportional to the support (LPP). Pies charts on branches represent normalised quartet score (QS) values for each alternative topology (blue = species tree topology (t1); green = first alternative topology; yellow = second alternative topology).

The concatenated plastid data matrix, spanning 57 samples and 67 CDS, also yielded high support values (UFB/SH-aLRT > 90%). The MLE topology inferred from these data was largely congruent with the nuclear phylogeny but exhibited some discordance, primarily along internal nodes (Figures 2 and 3). Family Bonnetiaceae was sister to Clusiaceae with moderate support (UFB = 68.60% / SH-aLRT = 79%), and both were sister to a clade composed of Calophyllaceae sister to Podostemaceae plus Hypericaceae (UFB/SH-aLRT = 100%). Within Clusiaceae, conflict was detected regarding the placement of *Moronobea* Aubl., which was sister to *Clusia rosea* Jacq. in the plastid tree (UFB/SH-aLRT = 100%, Figure 3). Calophyllaceae exhibits the highest conflict between nuclear and plastid phylogenies (Figure 3), with six out of eleven analyzed genera (*Caraipa, Haploclathra, Kayea, Kielmeyera, Marila,* and *Neotatea*) changing their placement across topologies. These shifts were supported by high bootstrap values in most cases (> 90%; Figures 2 and 3). Within Hypericaceae, conflict pertains to the placement of the tribes Cratoxyleae, Hypericeae, and Vismieae. The MLE plastid tree recovered tribe Cratoxyleae as sister to a Hypericeae and Vismieae clade (Figures 2 and 3), while the MSC and MLE nuclear species trees recovered Hypericeae as sister to a Cratoxyleae and Vismieae clade, in all cases with maximum support. The nuclear and plastid topologies were identical for tribe Cratoxyleae, which was not the case for tribes Hypericeae and Vismieae (Figures 2 and 3). In the plastid MLE tree, within tribe Hypericeae, *Hypericum elodes* L. was sister to *H. galioides* Lam. (UFB = 43.80%, SH-aLRT = 78%, Figure 2). Meanwhile, within tribe Vismieae, conflict was detected regarding the placement of *Vismia* (UFB/SH-aLRT = 100%, Figures 2 and 3).

**Figure 2.**
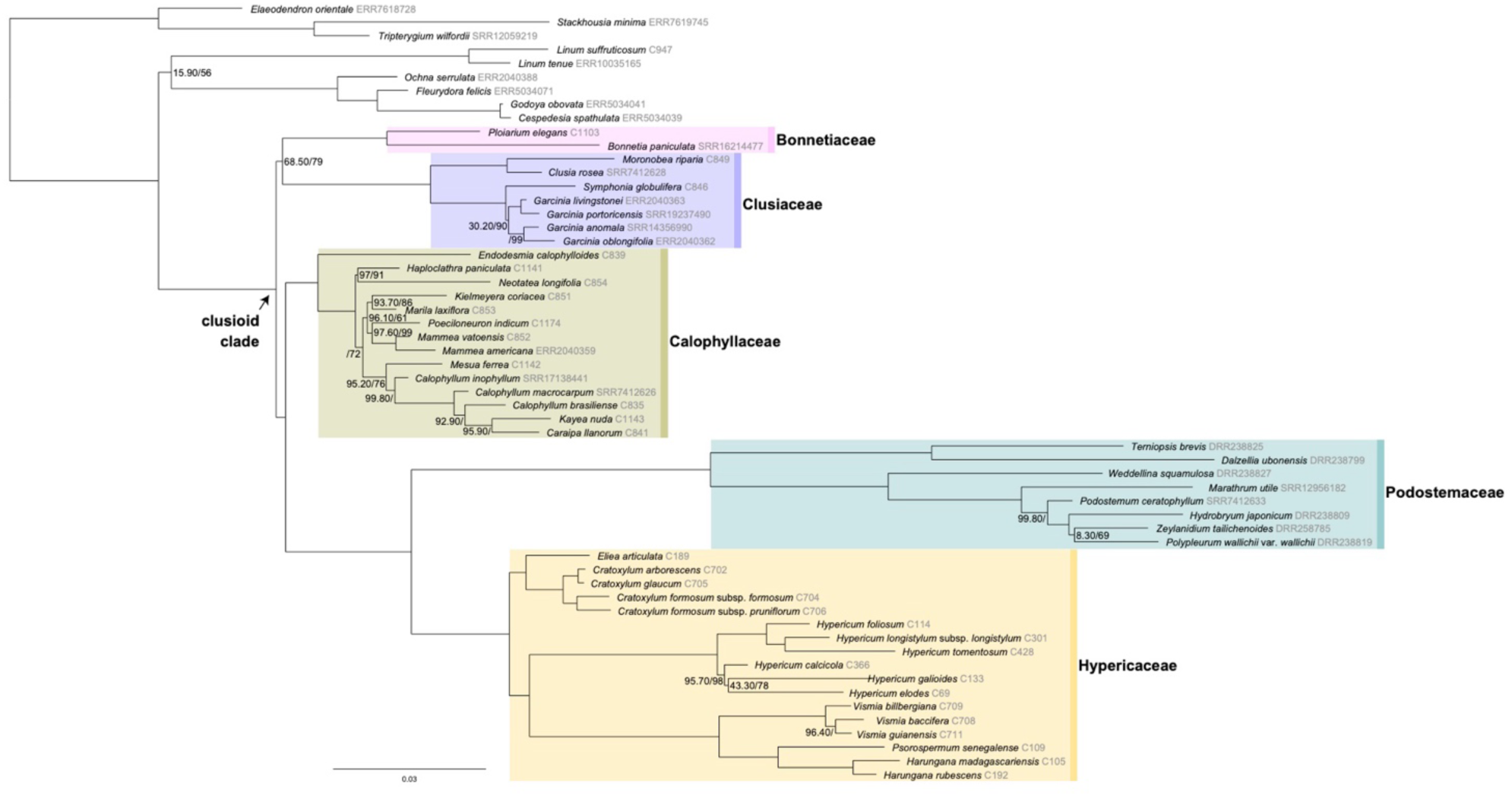
MLE nuclear phylogeny inferred in IQ-TREE2, for 57 clusioid-clade and related taxa, from 64 concatenated and partitioned plastid CDS mined with our custom amino acid plastid target file. Branch support shown for UFB and SH-aLRT if support value < 100.

**Figure 3.**
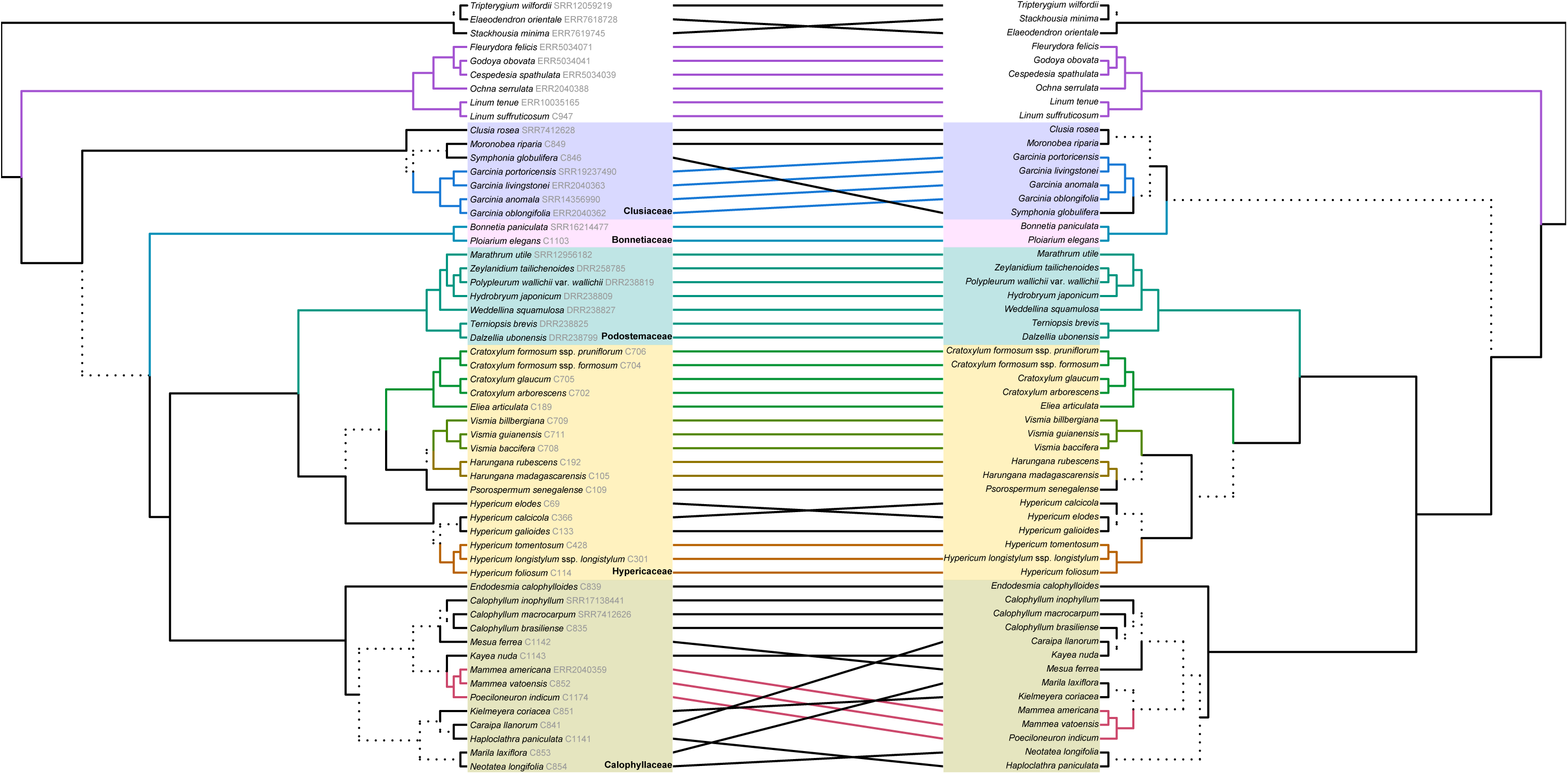
Tanglegram of the clusioid clade and related taxa comparing ultrametric topologies derived from the MSC nuclear (left) and MLE plastid (right) trees. Dashed lines highlight a combination of tips just present in one topology (i.e., unique clades). Subtrees shared across topologies are similarly colored.

### 3.6 Reticulation: Splits and Phylogenetic Networks

The SplitsTree network (Figure 4) resolved the five clusioid families into five distinct, well-supported groups (most bootstrap support values >80%), consistent with the phylogenetic trees. This splits network showed parallel edges at deep branches across all families, suggesting ancestral reticulation. Calophyllaceae and Podostemaceae exhibited the highest density of parallel edges at their stem. Clusiaceae displayed the longest parallel edges at its crown, reflecting extensive reticulation, and Bonnetiaceae showed the fewest parallel edges. Tip branches were longest in Podostemaceae.

**Figure 4.**
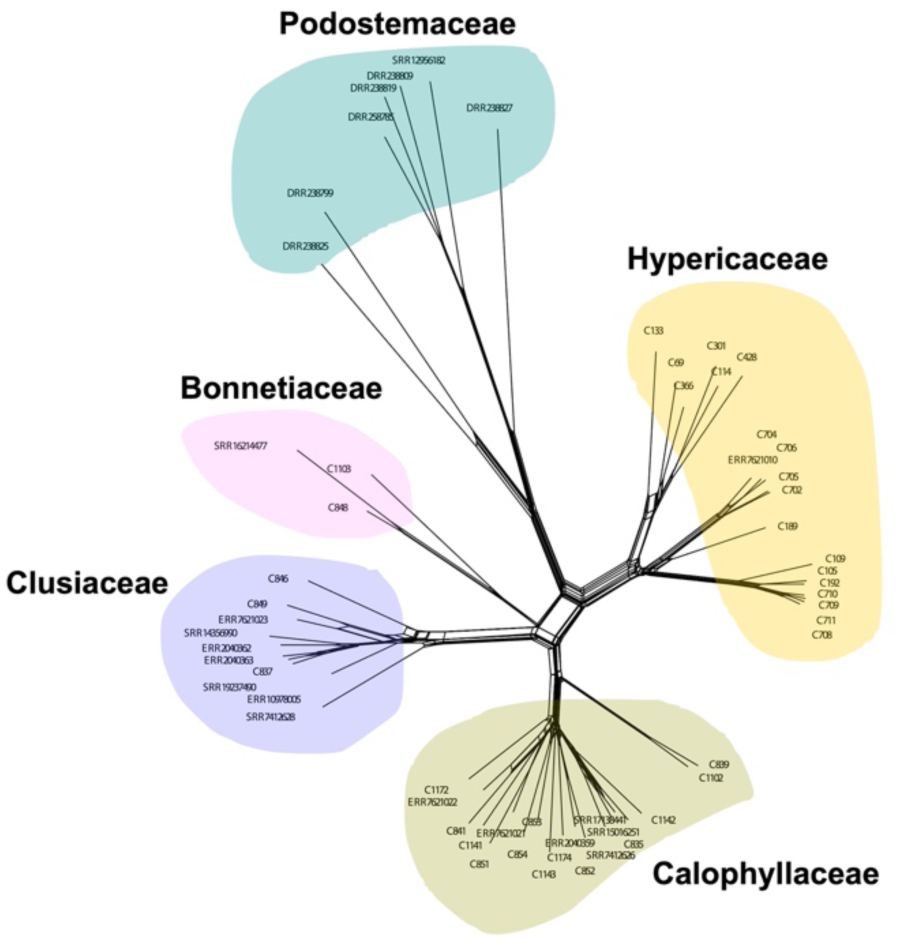
Nuclear splits network of the clusioid clade, inferred in SplitsTree (using the Neighbor-Net method, with uncorrected P-distances) from 552 concatenated, partitioned genes. Coloring scheme for families as in Figures 1 and 2.

The PhyloNet analysis recovered a topology similar to the MSC species tree (Figure S8). The best-fit phylogenetic network (log likelihood = −379.16) had four reticulations. This network indicates possible gene flow between the ancestor of *Linum suffruticosum* and an unsampled/extinct lineage ancestral to Podostemaceae. Within Podostemaceae, *Marathrum utile* Tul. and *Terniopsis brevis* M.Kato could have resulted from hybridization between unsampled/extinct lineages ancestral to both Podostemaceae and Hypericaceae. *Ploiarium elegans* Korth. (Bonnetiaceae) could have resulted from hybridization between unsampled/ancestral lineages to *Bonnetia sessilis* and family Calophyllaceae.

## 4 Discussion

Hyb-Seq has revolutionised the study of plant systematics (Dodsworth et al. 2019) by providing high-resolution data at multiple evolutionary scales and from DNA material of varying quality (Zuntini et al. 2024; Balant et al. 2025). The latest Hyb-Seq advancement pertains the integration of phylogenetically-broad enrichment panels (e.g., Angiosperms353, Johnson et al. 2019) with lineage-specific probe sets (e.g., Fonseca et al. 2023, 2024), enabling the development of custom kits that offer high resolution within groups, while retaining broad applicability across lineages.

### 4.1 The Clusioids626 Probe Set

Here we introduce the development and implementation of the Clusioids626 probe set, a novel tool that combines universal and custom probes to recover up to 626 nuclear ortholog genes chosen to resolve phylogenetic relationships across the clusioid clade. This kit is tailored to capture genetic data from all clusioid families (Bonnetiaceae, Calophyllaceae, Clusiaceae, Hypericaceae, and Podostemaceae), while also incorporating the universal Angiosperms353 targets. Unlike most existing combined kits, which are typically designed for the family-level or below (Bentz and Leebens-Mack 2024; Eserman et al. 2021; Fonseca et al. 2023, 2024; Musker et al. 2025; Ogutcen et al. 2021), the *Clusioids626* kit is the only one developed for the supra-familial level. This strategy removes the need for separate kits for each clusioid family, while maintaining the high resolution of lineage-specific probe sets and the broad applicability of universal enrichment panels. *Clusioids626* is also among the few probe sets specifically designed for malpighean lineages. To date, three other custom kits have been designed within this megadiverse order, targeting families Euphorbiaceae (431 loci, Villaverde et al. 2018), Ochnaceae (275 loci, Schneider et al. 2021; Shah et al. 2022), and Salicaceae (972 loci, Sanderson et al. 2020), and none of them incorporate the general-purpose Angiosperms353 targets.

Probe sets combining Angiosperms353 with custom targets have, on average, lower percentages of on-target reads (e.g., 30% for the Asparagaceae1726 kit, Bentz and Leebens-Mack 2024) than custom probe sets without universal targets (e.g., 86.7% for the Salicaceae972 kit, Sanderson et al. 2020). This pattern seems to reflect a trade-off between achieving broad taxonomic applicability at the cost of slightly reduced hybridization efficiency for certain taxa, which has been reported at supra-familial levels (Liu et al. 2019). However, samples processed with our Clusioids626 probe set yielded on average 50.4% on-target reads, which is comparable to the average capture efficiency detected for combined kits (e.g., 49.5% for the Annonaceae799 kit, Fonseca et al. 2024), but also for some specific kits within Malpighiales, such as the Euphorbiaceae431 kit (48.6%, Villaverde et al. 2018), while not as high as the average for the Salicaceae972 specific kit (see above). Differences in target recovery across studies may also be due to variations in molecular protocols (e.g., DNA purification, genomic library preparation, pooling strategies, hybridization settings) and computational workflows (e.g., mapping approach, target file sequence composition; Andermann et al. 2020).

Given our on-target recovery percent (50.4%), our probe set successfully captures a broad range of nuclear orthologs, providing sufficient phylogenetic signal for robust macroevolutionary analyses across the clusioid clade (see below). Its applicability at intraspecific levels remains to be explored, but we expect the Clusioids626 probe set to outmatch the Angiosperms353 enrichment panel at the microevolutionary scale, as it contains most Angiosperms353 targets plus 273 clusioid-specific ones. Beyond clusioid lineages, we were also able to mine most of our Clusioids626 targets (using our nuclear target file) for other malpighialean lineages (e.g., Linaceae, Ochnaceae) and even for taxa outside the order (i.e., three Celastrales; Table 1). This suggests that the Clusioids626 probe set is an efficient and adaptable tool for phylogenomics and, putatively, for population genomics research, not only within the clusioid clade but also across closely-related angiosperm groups. Clusioids626 thus offers a distinct advantage: by operating at an intermediate taxonomic level and combining universal and specific probes, it balances efficiency, specificity, and broad applicability. Furthermore, off-target reads provide valuable genetic information (i.e., organellar DNA) to further support evolutionary inference for clusioid Malpighiales (see below).

### 4.2 Clusioids626 vs. Angiosperms353

We show here that a combined kit, mixing universal and specific probes, offers additional advantages over universal kits by themselves. The custom Clusioids626 probe set outperforms the universal Angiosperms353 enrichment panel in gene recovery and capture success for the clusioid clade (Tables S6 and S8). This is consistent with other studies showing that lineage-specific probe sets improve locus recovery by targeting orthologs with higher affinity in a given clade (Ufimov et al. 2021). The inclusion of universal Angiosperms353 loci ensures compatibility with existing datasets, facilitating broader comparative analyses (Johnson et al. 2019; Baker et al. 2022; Zuntini et al. 2024), while the custom loci improve resolution for lineage-specific relationships by increasing variable sites (1,094.16 in Clusioids626 vs. 541.26 in Angiosperms353) and PICs (776.82 Clusioids626 vs. 423.90 Angiosperms353) median values. Despite comparable median values of P_PIC_ per aligned sequence length (0.36 for Clusioids626 vs. 0.44 for Angiosperms353), the Clusioids626 probe set resulted in more robust topological support values, with more nodes achieving maximum LPP (Table S10) and with a higher proportion of nodes resolving gene-tree conflict (QS > ⅓). Notably, both kits recovered a similar number of nodes achieving QS values > 0.75, indicating the power of both targeted sequencing kits to resolve phylogenetic relationships within the clusioid clade.

Previous studies (Chau et al 2018; Larridon et al. 2020; Siniscalchi et al. 2021; Fonseca et al. 2024) also suggest that universal loci (e.g., Angiosperms353) provide comparable phylogenetic resolution to taxon-specific targets, but highlight a critical advantage of custom kits: notably improved capture success and variable site recovery. This is also true for our Clusioids626 kit. Our results evidence that custom probe sets improve the quantity and quality of data, enabling more robust phylogenomic (and putatively population genomic) analyses by increasing the number of informative loci (and maybe even SNPs). Thus, while universal kits remain valuable for broad comparisons, custom optimizations offer a strategic advantage for clade-specific studies.

### 4.3 Clusioid Clade Phylogenomics

The Clusioids626 probe set was applied to resolve phylogenetic relationships among and within families in the clusioid clade. For the first time here, we place in a phylogeny eight clusioid species for which there was no sequence data available. Furthermore, we generate additional data for 36 clusioids and one Linaceae species. Relationships inferred in this study, for 58 species representing all five clusioid families, all eight clusioid tribes, and 34 of the 92 genera currently recognized in the group, are mostly resolved and strongly supported, both for the nuclear (Figures 1, S5 and S6) and plastid (Figures 2 and 3) trees.

We confirm, in both the nuclear and plastid topologies, the monophyly of the clusioid clade with maximum support (Cai et al. 2019, 2021; Ruhfel et al. 2011; Zuntini et al. 2024). Within the cusioid clade, our nuclear and plastid trees both recover Calophyllaceae as sister to Hypericaceae plus Podostemaceae, in line with previous findings (Cai et al. 2019; Ramírez-Barahona et al. 2020; Ruhfel et al. 2011; Sun et al. 2016; Xi et al. 2012). The placement of Podostemaceae within the clusioid clade has been recently disputed by Baker et al. (2022) in their order-level analysis of angiosperms, placing Podostemaceae outside Malpighiales, perhaps as a result of a long-branch attraction (LBA) artefact in their massive data matrix (but see Zuntini et al. 2024). It is quite remarkable that subtending long branches characterize the backbone of Podostemaceae in our study (Figures 1, 2, S5 and S6) and previous ones (Ramírez-Barahona et al. 2020; Li et al. 2021; although see Tippery et al. 2011; Philbrick et al. 2018; Bedoya et al. 2019). Meanwhile, branches within Podostemaceae subfamilies (Podostemoideae, Tristichoideae, and Weddellinoideae) are much shorter, suggesting fewer changes have accumulated since the most recent common ancestor (MRCA) of the sampled species diverged (Soltis et al. 2019). The longer branches leading to Podostemaceae compared to other clusioids could indicate faster rates of evolution in their plastomes, as suggested by Ruhfel et al. (2016) and Bedoya et al. (2019), which could be attributed to a faster life cycle and shorter generations (Smith and Donoghue 2008); Podostemaceae is mainly represented by annual herbs that rely on water levels to complete their life cycle. Other aquatic lineages, such as Hydrostachyaceae (order Cornales), exhibit long branches as well, which could also result in LBA artefacts (Fan and Xiang 2003; but see Thomas et al. 2021). Faster mutation rates have also been associated with life-history traits, such as plant height, genome size, and age at first reproduction (Lehtonen and Lanfear 2014; Bromham et al. 2015), and with pseudogenization following gene loss, that leads to a release of selective constraints (and hence faster mutation rates; Peredo et al. 2013; Li et al. 2023). The factors driving fast molecular evolution in Podostemaceae remain to be explored.

### 4.4 Cyto-Nuclear Discordance in the Clusioid Clade

Topological incongruence between genomic compartments in our study and previous ones, as expected, centres around placements that have posed problems for a long time and that remain unsolved (Figures 1–3 and S6). Specifically, the placement of families Bonnetiaceae and Clusiaceae, with respect to the remaining clusioid families. The backbone of our nuclear topology presents Clusiaceae as sister to all other clusioid families (Figure 1), in agreement with the results of Ramírez-Barahona et al. (2020). However, our plastid tree recovers Clusiaceae sister to Bonnetiaceae (Figures 2 and 3), in agreement with other studies based on plastome-level data (Xi et al. 2012; Trad et al. 2021) or on a few plastid and nuclear markers (Wurdack and Davis 2009; Ruhfel et al. 2011, 2016; Sun et al. 2016).

Some of the relationships identified for the nuclear compartment in our study contrast with those reported in previous studies relying on target capture sequencing. Specifically, Cai et al. (2021) placed Hypericaceae as sister to all other clusioid families and Clusiaceae as sister to a Bonnetiaceae and Calophyllaceae clade, based on 423 ultra-conserved-like elements (ULEs; Buddenhagen et al. 2016). It should be noted that the ultra-conserved-like nature of these targets limits the phylogenetic signal they might present for phylogenetic inference (vital for complex plant lineages, such as the malpighialean angiosperms). Nuclear species trees inferred from Angiosperms353 targets (Baker et al. 2022; Zuntini et al. 2024) have previously placed Bonnetiaceae as sister to the remaining clusioid families with full support. Apart from methodological aspects potentially explaining topological incongruences between our study and previous ones, the degree of conflict revealed for clusioid clade families might stem from biological processes such as ILS, duplication (with or without loss), and/or hybridization (Larson et al. 2024), suggesting that the phylogenetic reconstruction of the clusioid clade is a major challenge, in line with the difficulties encountered in the reconstruction of Malpighiales as a whole (Cai et al. 2021).

### 4.5 Reticulation in the Clusioid Clade

The overall concordance between nuclear MLE and MSC phylogenies underscores the robustness of our nuclear targets for resolving clusioid clade relationships. However, considerable cyto-nuclear discordance (between nuclear and plastid trees) highlights divergent evolutionary histories across genomic compartments, a phenomenon increasingly recognized in angiosperms (Stull et al. 2020, 2023; Kandziora et al. 2022; Joyce et al. 2025). The splits network (Figure 4), inferred with SplitsTree 6, revealed widespread parallel edges at deep nodes across all families, suggestive of ancestral phylogenetic conflict during their divergence, particularly pronounced in Podostemaceae and Calophyllaceae. Deep reticulation could result from several causes, such as ILS during rapid diversification leading to conflicting phylogenetic signal (Kandziora et al. 2022), ancient hybridization and reticulation due to ecological introgression (Nge et al. 2021), allopolyploidization (Soltis et al. 2019), ecological and biogeographic factors, and even methodological artefacts (Steenwyk et al.

2023; Bjornson et al. 2024; Joyce et al. 2025). In the clusioid clade, Podostemaceae exhibits the longest terminal branches and the densest reticulation at the stem, consistent with an accelerated molecular evolution scenario associated with adaptation to extreme aquatic habitats (Jin et al. 2020). Clusiaceae shows the longest parallel edges at its crown, suggesting recurrent gene flow, possibly driven by ecological versatility (e.g., facultative CAM photosynthesis, epiphytic growth; Qiu et al. 2023) and high species diversity (Cardoso et al. 2017), facilitating hybridization in neotropical niches. Bonnetiaceae exhibits minimal reticulation, probably reflecting a stable evolutionary trajectory (Ruhfel et al. 2011). The phylogenetic network (Figure S8), inferred with PhyloNet, identifies gene flow between ancestral Linaceae and Podostemaceae, as well as between Hypericaceae and Podostemaceae. These results align with the deep reticulations and parallel edges of the splits network, suggesting historical introgression or hybridization during the clusioid radiation. The coexistence of nuclear concordance and plastid discordance emphasizes the need for integrative approaches combining universal and taxon-specific loci. While universal kits (Angiosperms353) anchor deep relationships, combined kits such as Clusioids626 improve the resolution at shallow evolutionary levels.

## Supporting information

Suppl_mat_Tables_S1-S12

## Acknowledgements

We would like to thank Mario Rincón-Barrado for bioinformatic resources and Mónica García-Gallo for technical support. The authors are very thankful to the following herbaria for providing plant material: A, AAU, BM, F, K, MA, MO, NY, and U for helping with loan of specimens. I.V.-M., I.M-I., and A.S. were supported by the Atracción de Talento CAM program (2019-T1/AMB-12648) and by project PID2020-120145GA-I00 funded by MICIU/AEI /10.13039/501100011033. I.V.-M. is supported by a Fundación Ramón Areces postdoctoral research fellowship (Ref.: BEVP35A7117). L.P. benefited from a Ramón y Cajal grant (Ref.: RYC2021-034942-I) funded by MCIN/AEI/10.13039/501100011033 and by European Union “NextGenerationEU”/PRTR. This research project was made possible through access granted by the Galician Supercomputing Center (CESGA) to its supercomputing infrastructure. The supercomputer FinisTerrae III and its permanent data storage system have been funded by the Spanish Ministry of Science and Innovation, the Galician Government, and the European Regional Development Fund (ERDF).

## Data Accessibility Statement

The raw reads are available in the NCBI Sequence Read Archive (accession no. PRJNA1284379). The intermediate files (trimmed alignments, gene and species trees, and matrices used for phylogenetic networks), the scripts, and the workflow can be found on figshare (https://figshare.com/s/052464d25042e82170f2).

## Author Contributions

A.S.M., L.P., and I.V.-M. conceived the study; A.S.M., I.M.-I., and N.N. obtained the samples. I.V.-M. performed bioinformatic analyses to identify the clusioid targets. I.V.-M., I.M.-I., and L.P. did molecular work and analyzed the data; I.V.-M., I.M.-I., L.P., and A.S.M. wrote the manuscript, with feedback from N.N. A.S.M. provided funding. All authors approved the final version of the manuscript.

