## Supplementary material for "Enrichment Probe Sets Combining Universal and Lineage-Specific Targets Help Resolve Recalcitrant Lineages": Suppl_figures

### Table of Contents:


| | |
|---|---|
| **Figure S1** | Page 2 |
| **Figure S2** | Page 3 |
| **Figure S3** | Page 4 |
| **Figure S4** | Page 5 |
| **Figure S5** | Page 6 |
| **Figure S6** | Page 7 |
| **Figure S7** | Page 8 |
| **Figure S8** | Page 9 |

### Supplementary Figures

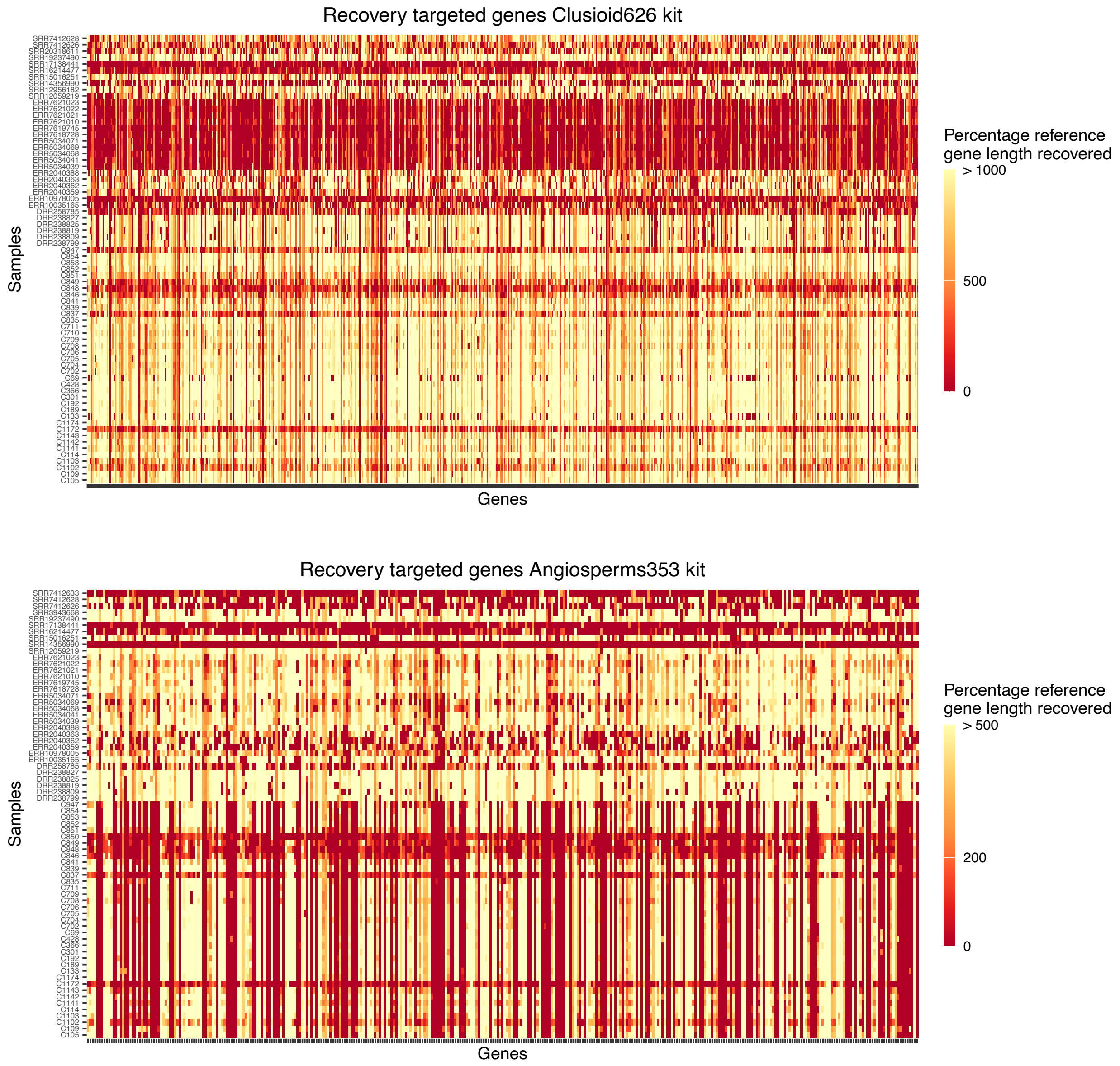


**Figure S1.** Heatmap of the recovered sequence length of (A) the 626 genes targeted with the Clusioids626 probe set, and (B) the 353 genes targeted with Angiosperms353 enrichment panel.

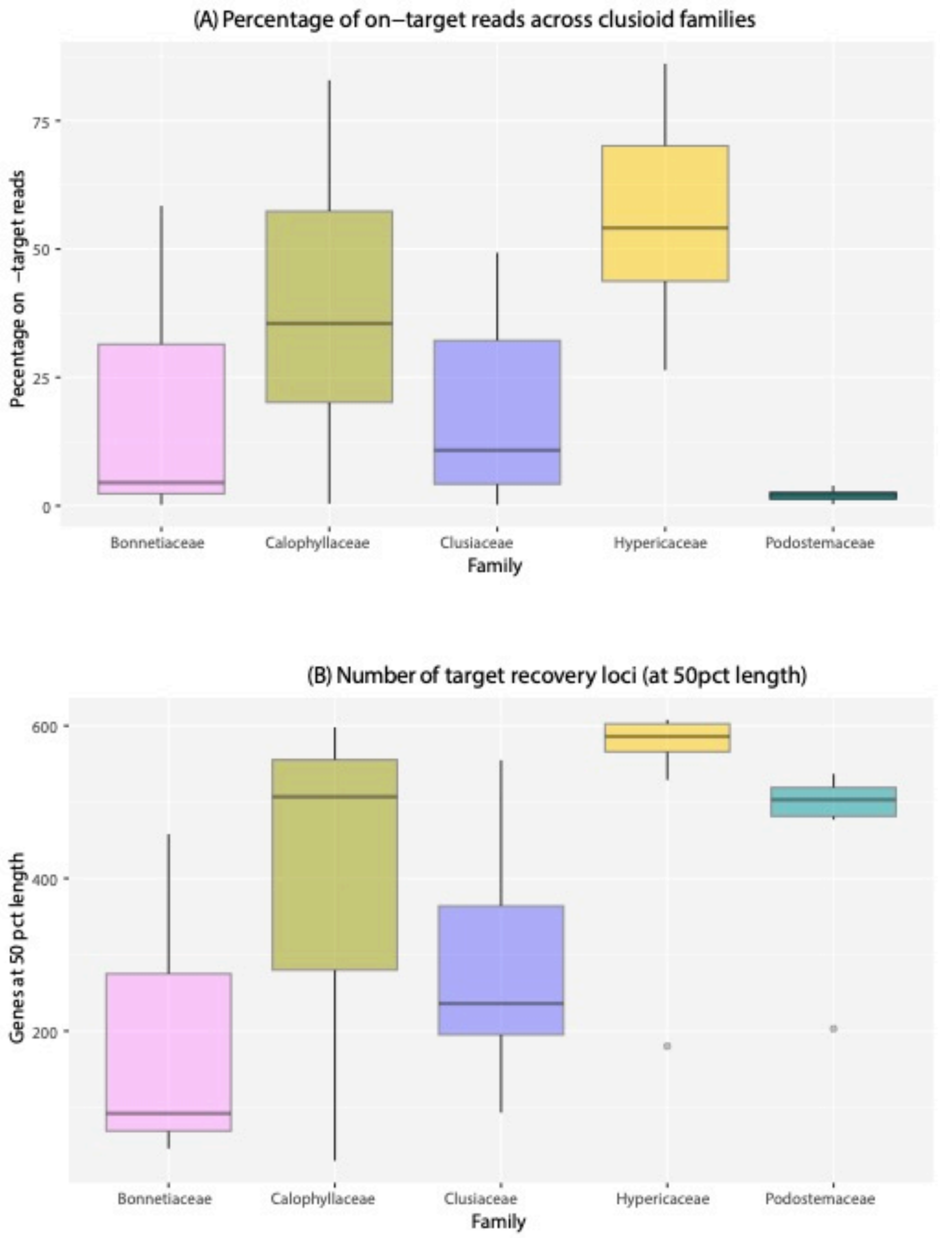


**Figure S2.** Boxplots showing (A) the percentage of on-target reads and (B) the total number of Clusioids626 target loci recovered at a 50% target length for each clusioid family.

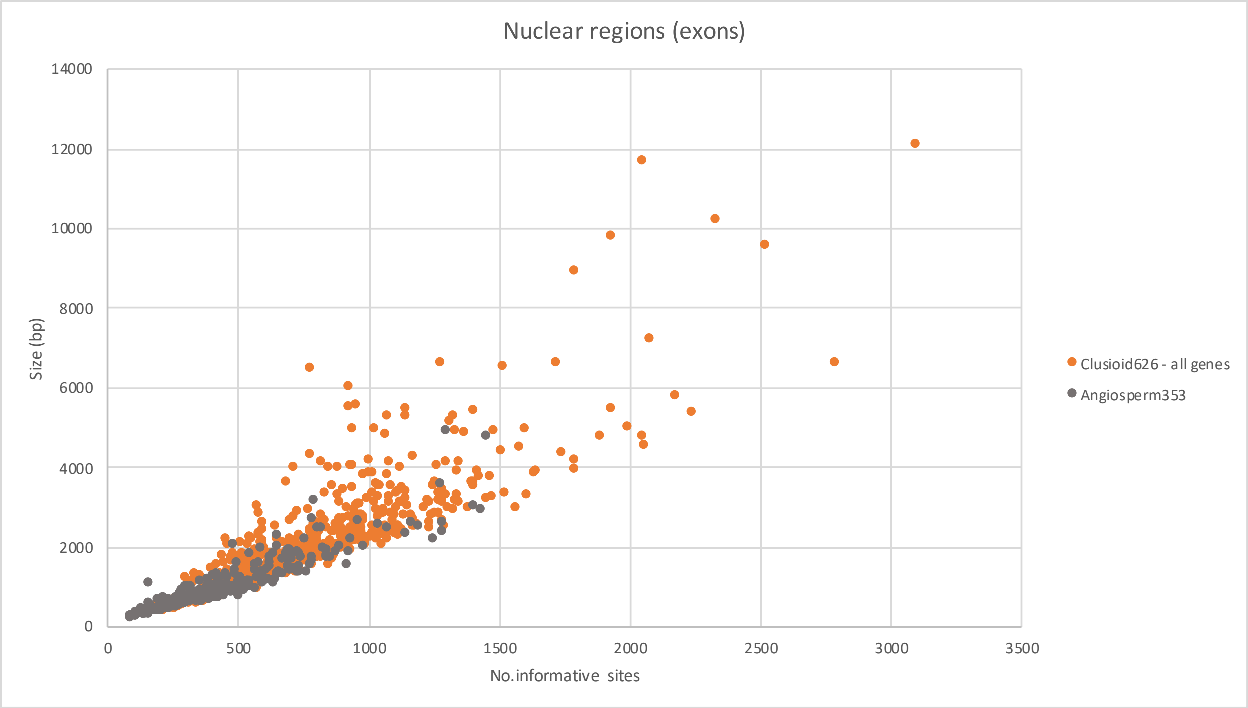


**Figure S3.** Scatter plot showing the relationship between the size and number of Parsimony-informative characters (PICs) for each of the 608 genes from the Clusioids626 probe set and 345 genes from the Angiosperms353 enrichment panel.

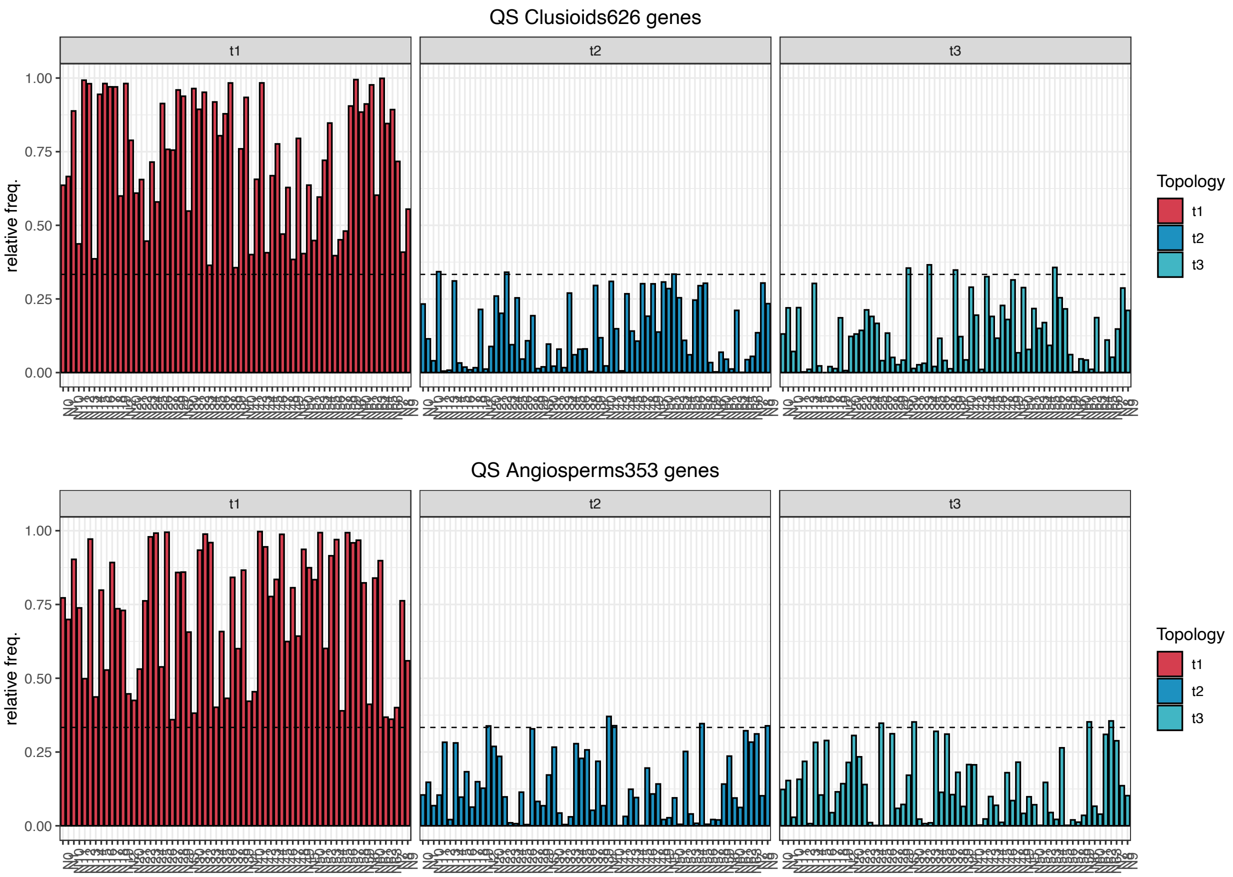


**Figure S4.** Quartet score (QS) frequencies for focal internal nodes of the ASTRAL species tree, inferred from shrunk, trimmed gene trees. (A) QS frequencies for the 67 nodes of the species tree obtained using the Clusioids626 target file. (B) QS frequencies for the 63 nodes of the species tree obtained using the Angiosperms353 target file. The main topology (t1) is shown in red, and the two alternative topologies (t2, t3) are shown in blue. Dotted lines indicate the 1/3 threshold.

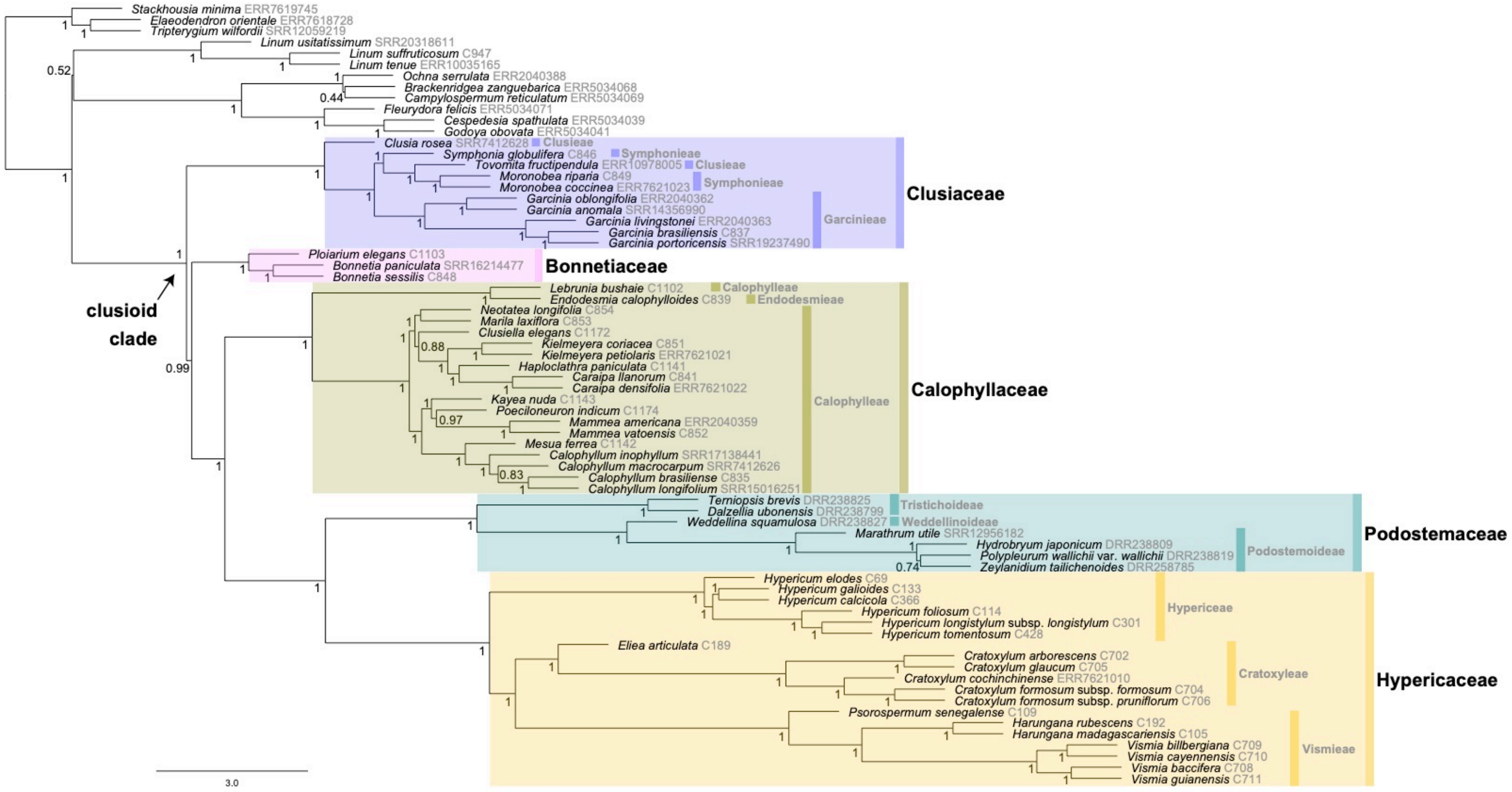


**Figure S5**. Coalescent-based species trees generated by ASTRAL-III for 70 samples of clusioid clade and related taxa enriched with Clusioids626, based on 552 exon loci. Node values represent LPP for the main topology and are equal to one unless noted otherwise.

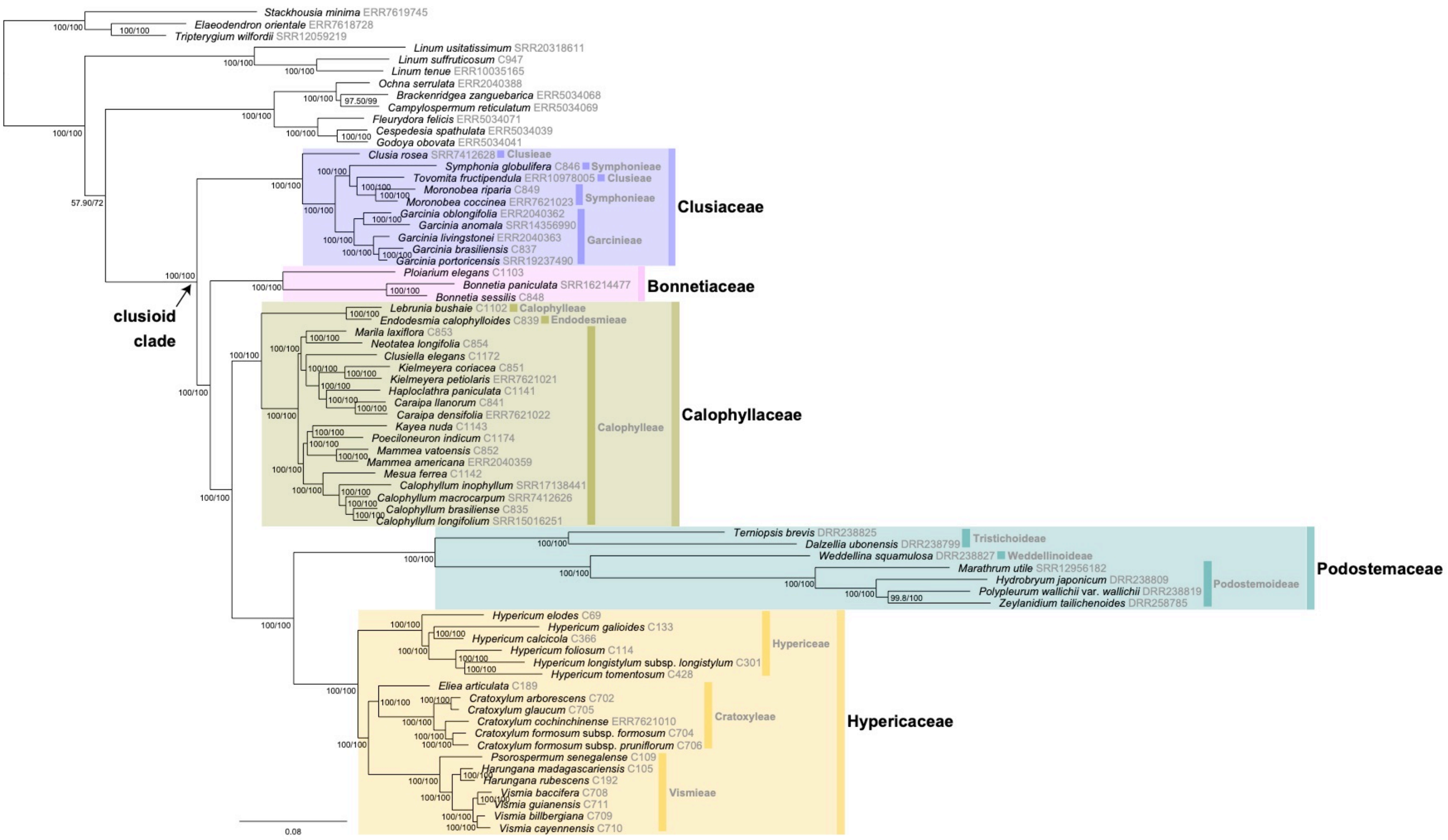


**Figure S6**. Maximum likelihood estimation-based phylogenetic reconstruction of the clusioid clade inferred using 552 loci concatenated and mined with the Clusioids626 target file. Values at the nodes represent bootstrap support and Shimodaira-Hasegawa-like approximate likelihood ratio tests (SH-aLRT) values, respectively.

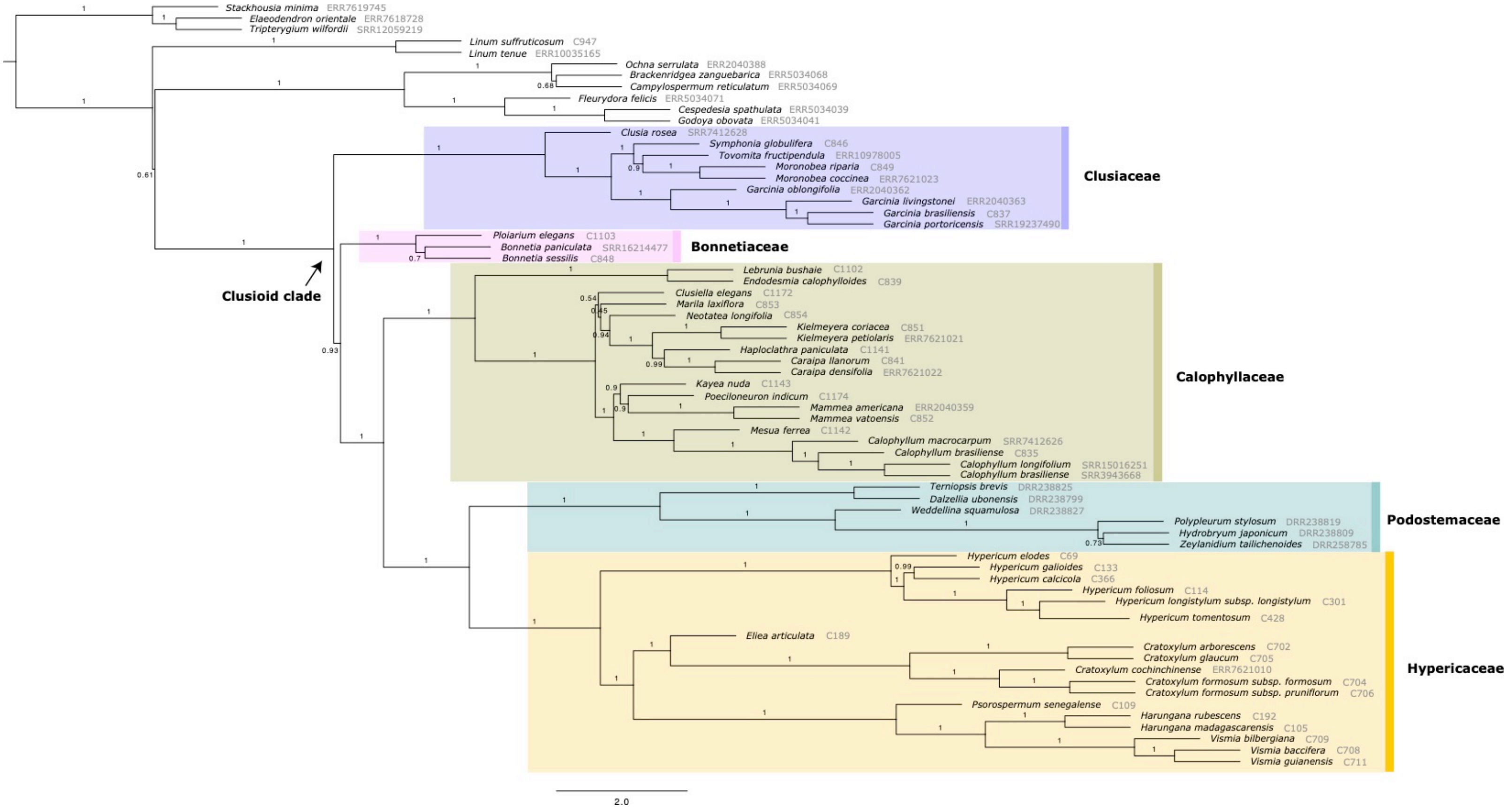


**Figure S7.** Phylogenetic reconstructions using ASTRAL of the relationships in the clusioid clade and related taxa for 66 accessions from 337 loci mined with the Angiosperms353 target file. Branch values represent LPP values for the main topology.

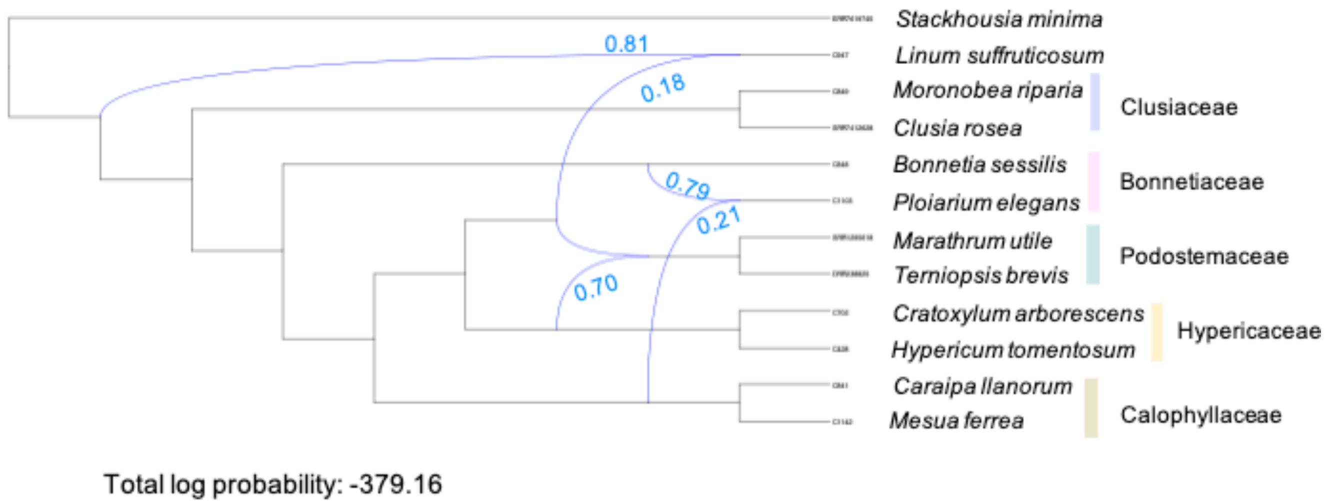


**Figure S8**. PhyloNet estimation results for the simplified phylogenetic tree of the clusioid clade inferred with the InferNetwork_MPL method, based on 63 rooted nuclear gene trees. The blue edges denote the reticulation events identified with numbers next to the edges, denoting the inheritance probabilities.
